## Supplementary Figures for "A virus-encoded protein suppresses methylation of the viral genome in the Cajal body through its interaction with AGO4"

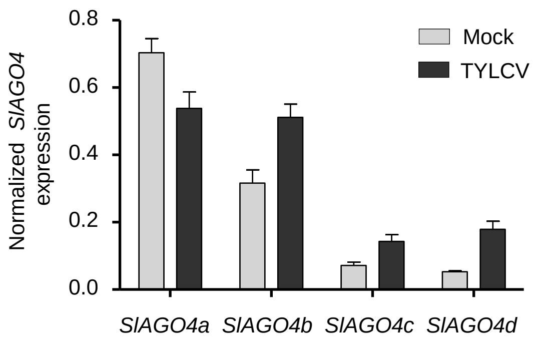


**Figure S1. *SlAGO4* expression in TYLCV-infected and control tomato plants.** *SlAGO4a/b/c/d* expression in TYLCV-infected or control (mock-inoculated) tomato plants at 3 weeks post-inoculation (wpi), as measured by qRT-PCR. Gene expression was normalized to *SlActin.* Values are the mean of three independent biological replicates; error bars indicate SEM.

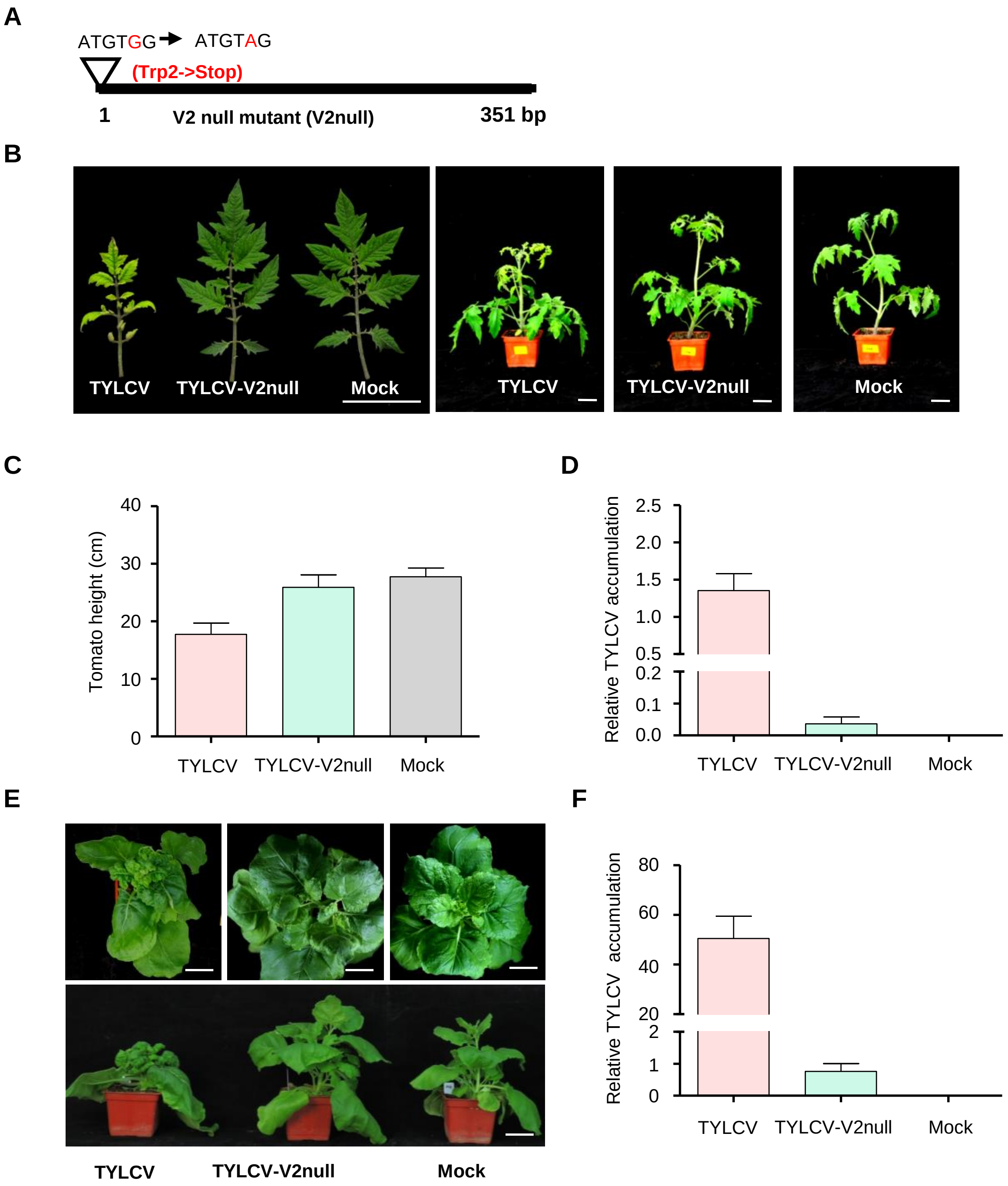


**Figure S2. V2 is essential for systemic infection in tomato and *N. benthamiana*.**

(A) Design of the V2 null TYLCV mutant used in this work. The second codon of the V2 ORF, originally encoding a Trp (TGG), is mutated to STOP codon (TAG).

(B) Representative pictures of tomato plants infected with TYLCV WT or V2 null mutant (TYLCV-V2null) or mock-inoculated. Photographs were taken at 3 weeks post-inoculation (wpi). Bar, 5cm.

(C) Height of tomato plants infected with TYLCV WT or V2 null mutant (TYLCV-V2null) or mock-inoculated at 3 wpi. Values are the mean of five independent biological replicates; error bars indicate SEM.

(D) Viral (TYLCV) accumulation in tomato plants infected with TYLCV WT or V2 null mutant (TYLCV-V2null) or mock-inoculated at 3 wpi, measured by qPCR. Each sample corresponds to the apical leaves from six plants. The accumulation of viral DNA is normalized to the *25S ribosomal RNA interspacer* (*ITS*). Values are the mean of six independent biological replicates; error bars indicate SEM.

(E) Representative pictures of *N. benthamiana* plants infected with TYLCV WT or V2 null mutant (TYLCV-V2null) or mock-inoculated. Photographs were taken at 3 weeks post-inoculation (wpi). Bar, 5cm.

(F) Viral (TYLCV) accumulation in *N. benthamiana* plants infected with TYLCV WT or V2 null mutant (TYLCV-V2null) or mock-inoculated at 3 wpi, measured by qPCR. Each sample corresponds to the apical leaves from six plants. The accumulation of viral DNA is normalized to the *25S ribosomal RNA interspacer (ITS)*. Values are the mean of six independent biological replicates; error bars indicate SEM.

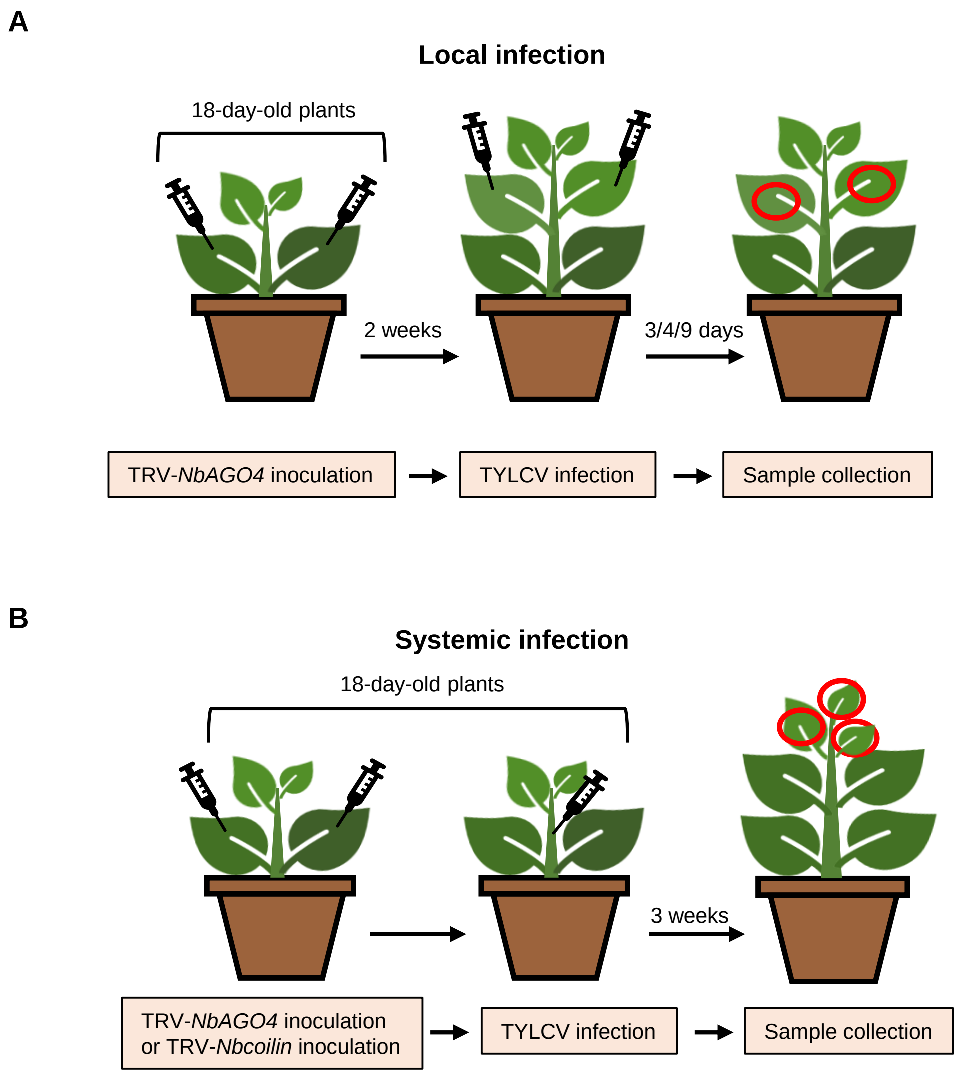


**Figure S3. Experimental design for local and systemic TYLCV infection assays in *NbAGO4*-silenced *N. benthamiana* plants.**

(A) Experimental design for local TYLCV infection assays. *A. tumefaciens* carrying the TRV-EV or TRV-*NbAGO4* infectious clones were inoculated on 18-day-old *N. benthamiana* cotyledons. 2 weeks later, young leaves were infiltrated with *A. tumefaciens* carrying the TYLCV infectious clone (WT or V2null) and leaf patches were collected at 3, 4, or 9 days post-inoculation (dpi).

(B) Experimental design for systemic TYLCV infection assays. *A. tumefaciens* carrying the TRV-EV or TRV-*NbAGO4* infectious clones were inoculated on 18-day-old *N. benthamiana* cotyledons. At the same time, *A. tumefaciens* carrying the TYLCV infectious clone (WT or V2null) were injected into plant stems. The top three leaves were collected at 3 weeks post-inoculation (wpi).

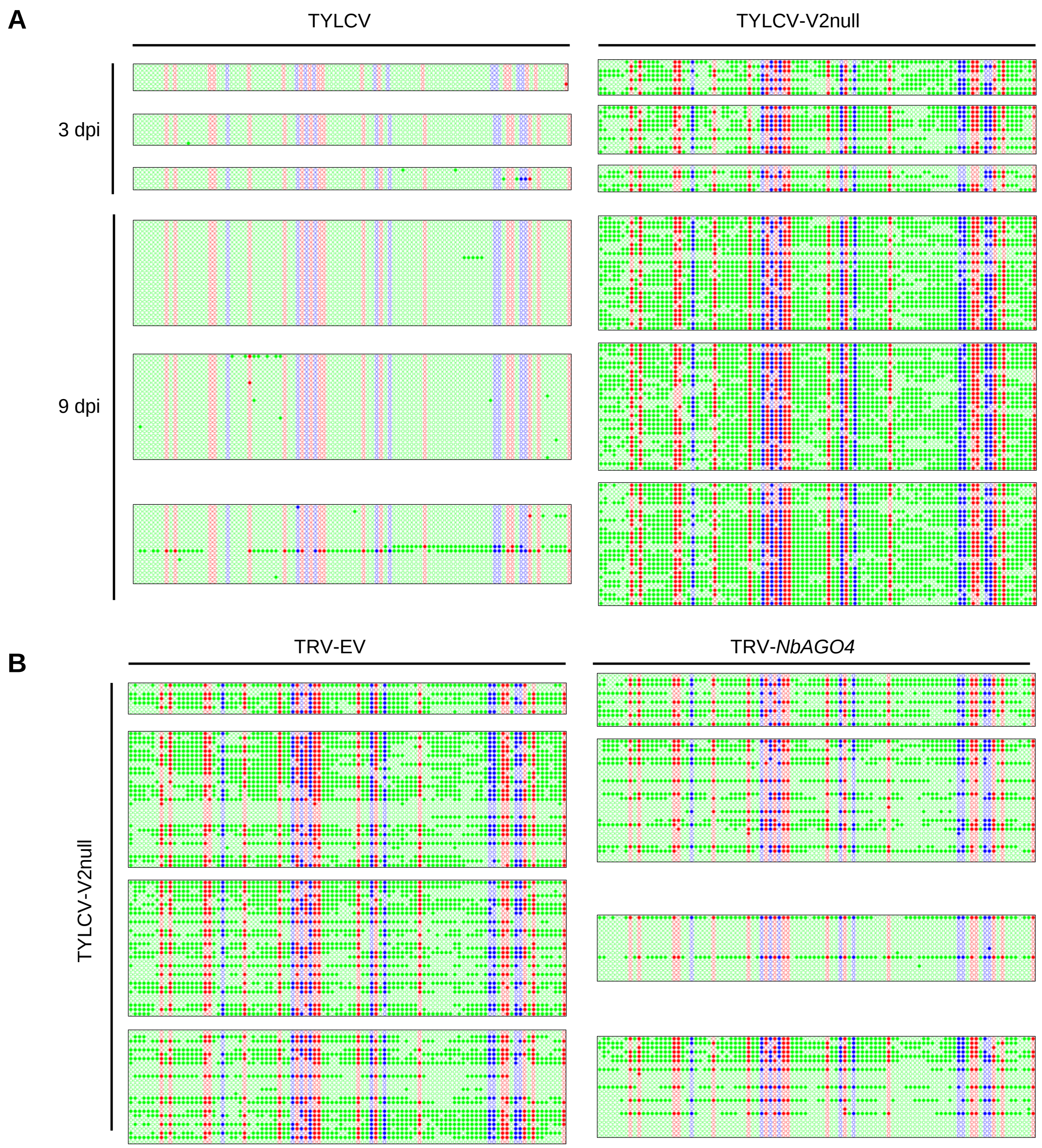


**Figure S4. Original single-base resolution bisulfite sequencing data of the intergenic region (IR) of TYLCV (WT and V2 null mutant) in local infection assays, related to Figure 4A, B.**

(A) Original single-base resolution bisulfite sequencing data for Figure 4A. At least 5 individual clones were sequenced per replicate and sample at 3 dpi, and >18 individual were sequenced per replicate and sample at 9 dpi. Each single circle, corresponding to a cytosine, is colored in blue, red, or green, representing the CHG, CG or CHH contexts, respectively. Methylated cytosines are represented by filled circles, while unmethylated cytosines are represented by empty circles.

(B) Original single-base resolution bisulfite sequencing data for Figure 4B. At least 7 individual clones were sequenced per sample at 4 dpi in the first replicate, and > 15 individual clones were sequenced per sample at 4dpi in replicates second to fourth. Each single circle, corresponding to a cytosine, is colored in blue, red, or green, representing the CHG, CG or CHH contexts, respectively. Methylated cytosines are represented by filled circles, while unmethylated cytosines are represented by empty circles.

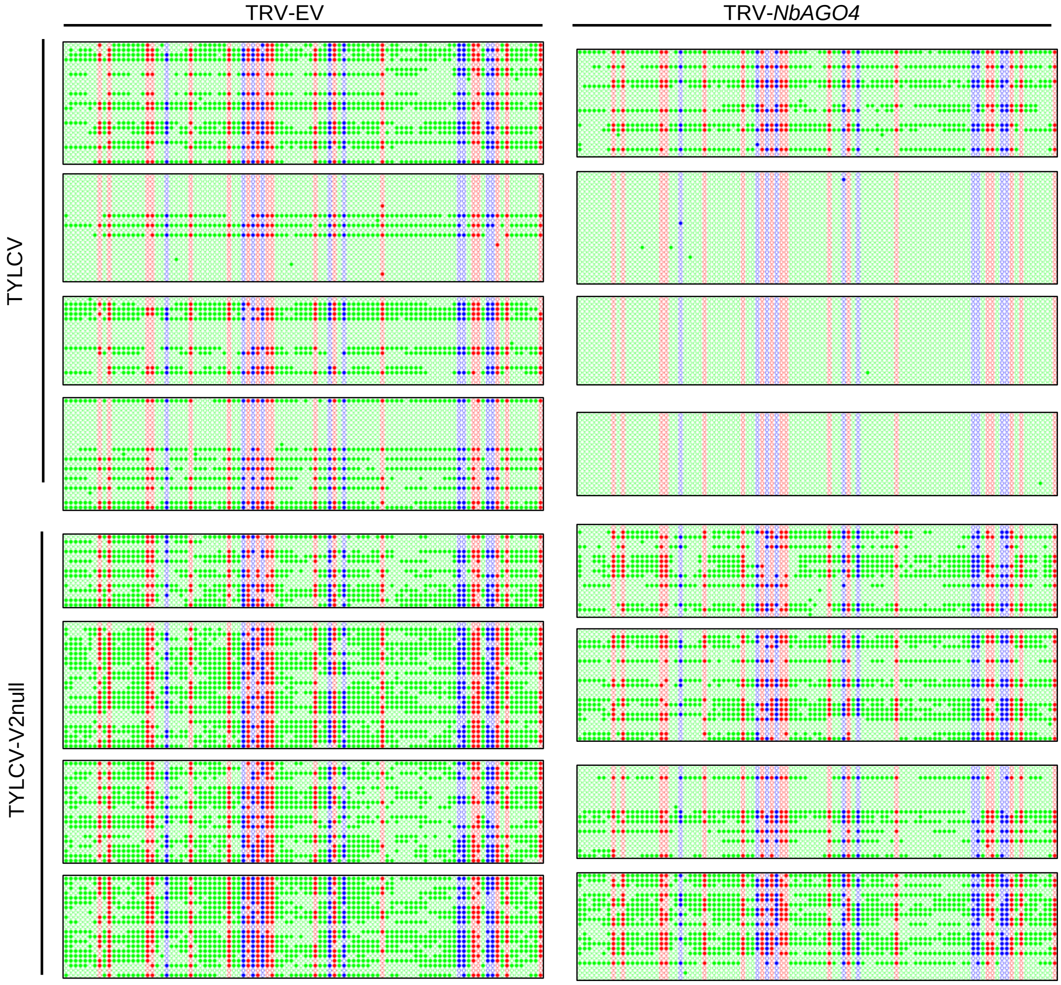


**Figure S5. Original single-base resolution bisulfite sequencing data of the intergenic region (IR) of TYLCV and TYLCV-V2null in systemic infection assays, related to Figures 4C.** Original single-base resolution bisulfite sequencing data for Figure 4C. >14 individual clones were sequenced per sample and replicate. Each single circle, corresponding to a cytosine, is colored in blue, red, or green, representing the CHG, CG or CHH contexts, respectively. Methylated cytosines are represented by filled circles, while unmethylated cytosines are represented by empty circles.

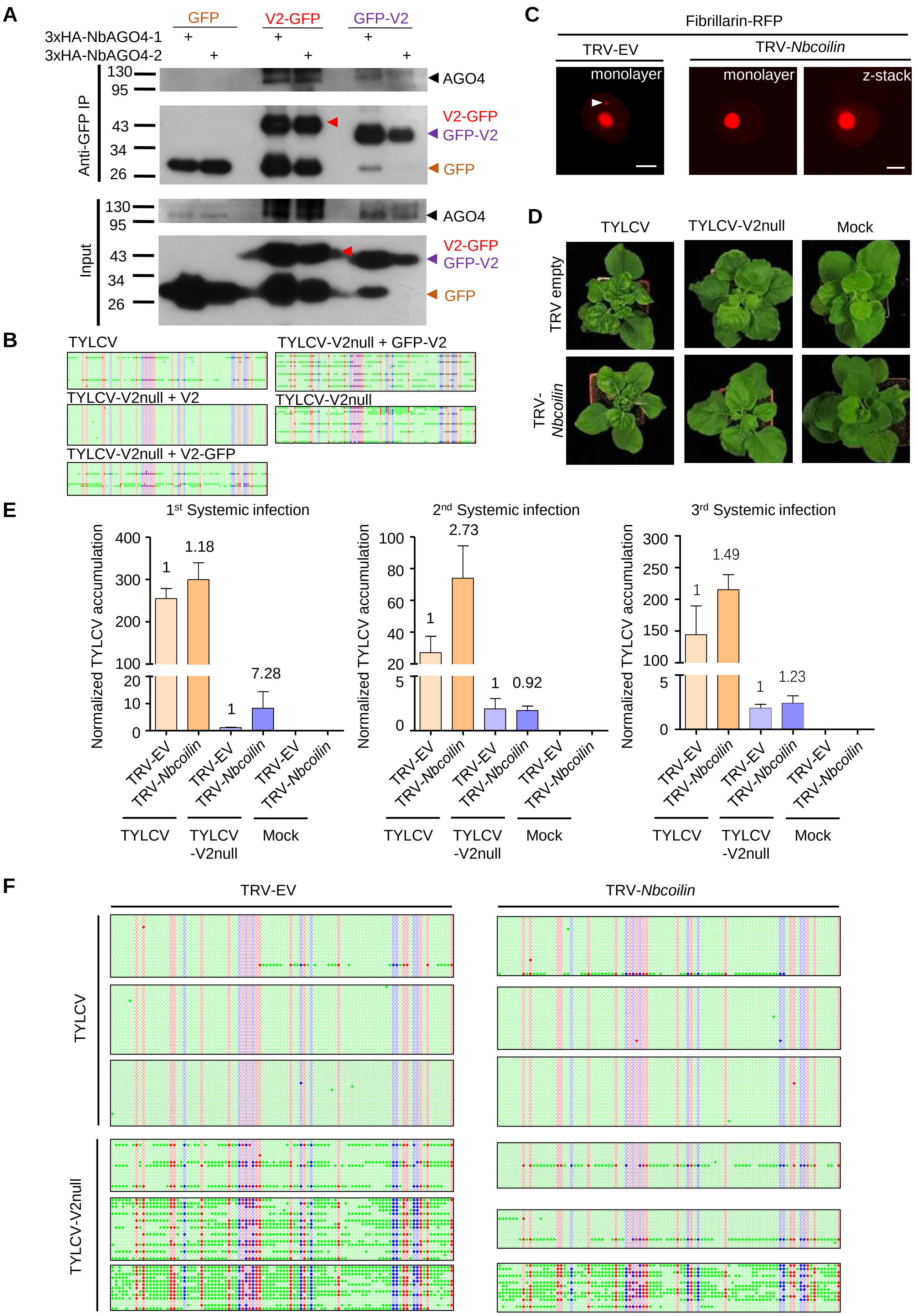


**Figure S6:** The relevance of the Cajal body localization of the V2-AGO4 interaction and TYLCV infection in *Nbcoilin*-silenced plants.

(A) 3xHA-NbAGO4-1 and 3xHA-NbAGO4-2 interact with V2-GFP and GFP-V2 in co-immunoprecipitation (co-IP) assays upon transient expression in *N. benthamiana*. Free GFP was used as negative control. Three independent biological replicates were performed with similar results.

(B) Original single-base resolution bisulfite sequencing data for Figure 7B. >16 individual clones were sequenced per sample and replicate. Each single circle, corresponding to a cytosine, is colored in blue, red, or green, representing the CHG, CG or CHH contexts, respectively. Methylated cytosines are represented by filled circles, while unmethylated cytosines are represented by empty circles.

(C) No Cajal body was observed in the nuclei of *Nbcoilin*-silenced plants, whereas in control plants at least one Cajal body was normally present in each nucleus; these results are in agreement Shaw et al., 2014. Fibrillarin-RFP, used as a nucleolus and Cajal body marker, was transiently expressed in *N. benthamiana* epidermal cells of *Nbcoilin*-silenced (TRV-*Nbcoilin*) and control plants (TRV-EV). Confocal images were taken at two days after infiltration. Arrowheads indicate the position of the Cajal body. Bar, 5μm. This experiment was repeated three times with same results.

(D) Representative pictures of *N. benthamiana* plants infected with the indicated combinations of viruses. Photographs were taken at 3 weeks post-inoculation (wpi).

(E) Viral (TYLCV) accumulation in systemic infections in *Nbcoilin*-silenced or control plants, measured by qPCR. Apical leaves from six plants were collected at 3 wpi. The experimental design is shown in Figure S3B. The accumulation of viral DNA is normalized to the *25S ribosomal RNA interspacer (ITS)*. Results from three independent experiments are shown. Values are the mean of six independent biological replicates; error bars indicate SEM. The relative fold change of viral accumulation between *Nbcoilin*-silenced plants and control plants is shown above each column.

(F) Original single-base resolution bisulfite sequencing data for Figure 6D. >11 individual clones were sequenced per sample and replicate. Each single circle, corresponding to a cytosine, is colored in blue, red, or green, representing the CHG, CG or CHH contexts, respectively. Methylated cytosines are represented by filled circles, while unmethylated cytosines are represented by empty circles.

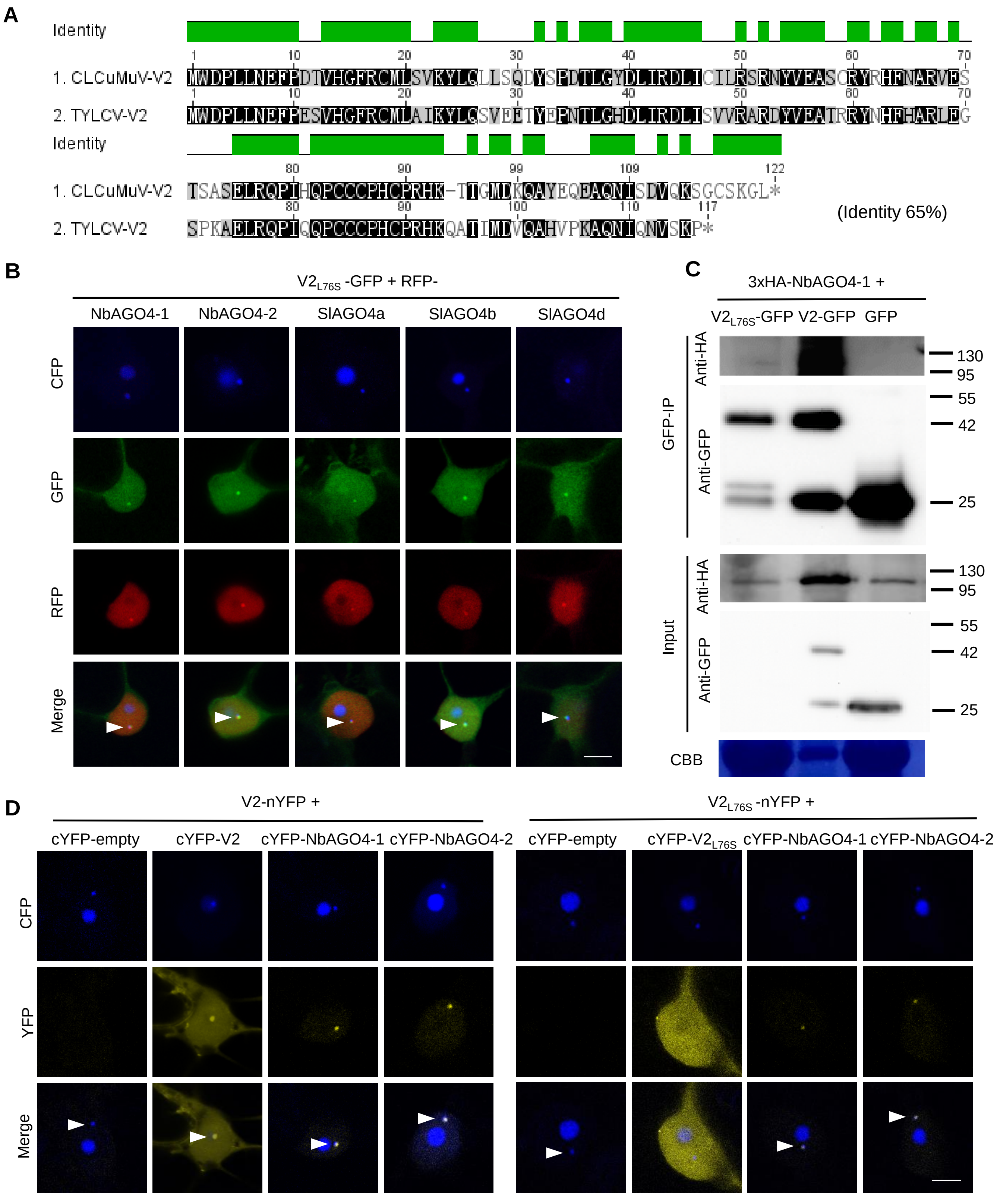


**Figure S7. TYLCV $V2_{L76S}$ interacts with AGO4 in the Cajal body.**

(A) Alignment of the amino acid sequences of V2 from *Cotton Leaf Curl Multan virus* (CLCuMuV) and V2 from TYLCV. The alignment was performed by Geneious (https://www.geneious.com). Black background indicates conservation. The identity of these two proteins is 65%.

(B) $V2_{L76S}$-GFP and RFP-AGO4 co-localize in the Cajal body. CFP-Fibrillarin, $V2_{L76S}$-GFP and RFP-NbAGO4-1/2 or RFP-SlAGO4a/b/d were transiently co-expressed in *N. benthamiana* epidermal cells. CFP-Fibrillarin is used as a nucleolus and Cajal body marker. Confocal images were taken at two days after infiltration. Arrowheads indicate the position of the Cajal body. Bar, 5μm. This experiment was repeated three times with similar results.

(C) 3xHA-NbAGO4-1 interacts with V2-GFP and $V2_{L76S}$-GFP in co-immunoprecipitation (co-IP) assays upon transient expression in *N. benthamiana*. Free GFP was used as negative control. CBB, Coomassie brilliant blue staining. The V2-GFP sample was diluted 1/20 for western blot to reach a protein amount comparable to that of $V2_{L76S}$-GFP. Three independent biological replicates were performed with similar results.

(D) $V2_{L76S}$ interacts with AGO4 in the Cajal body. The N-terminal half of the YFP fused to $V2_{L76S}$ ($V2_{L76S}$-nYFP) was transiently co-expressed with the C-terminal half of the YFP alone (cYFP, as a negative control), or cYFP-NbAGO4, cYFP-SlAGO4, or cYFP- $V2_{L76S}$ in *N. benthamiana* leaves. CFP-Fibrillarin was used as a nucleolus and Cajal body marker. V2 was used as a control. Yellow fluorescence indicates a positive interaction. Arrowheads indicate the position of the Cajal body. Bar, 5μm. This experiment was repeated three times with similar results.

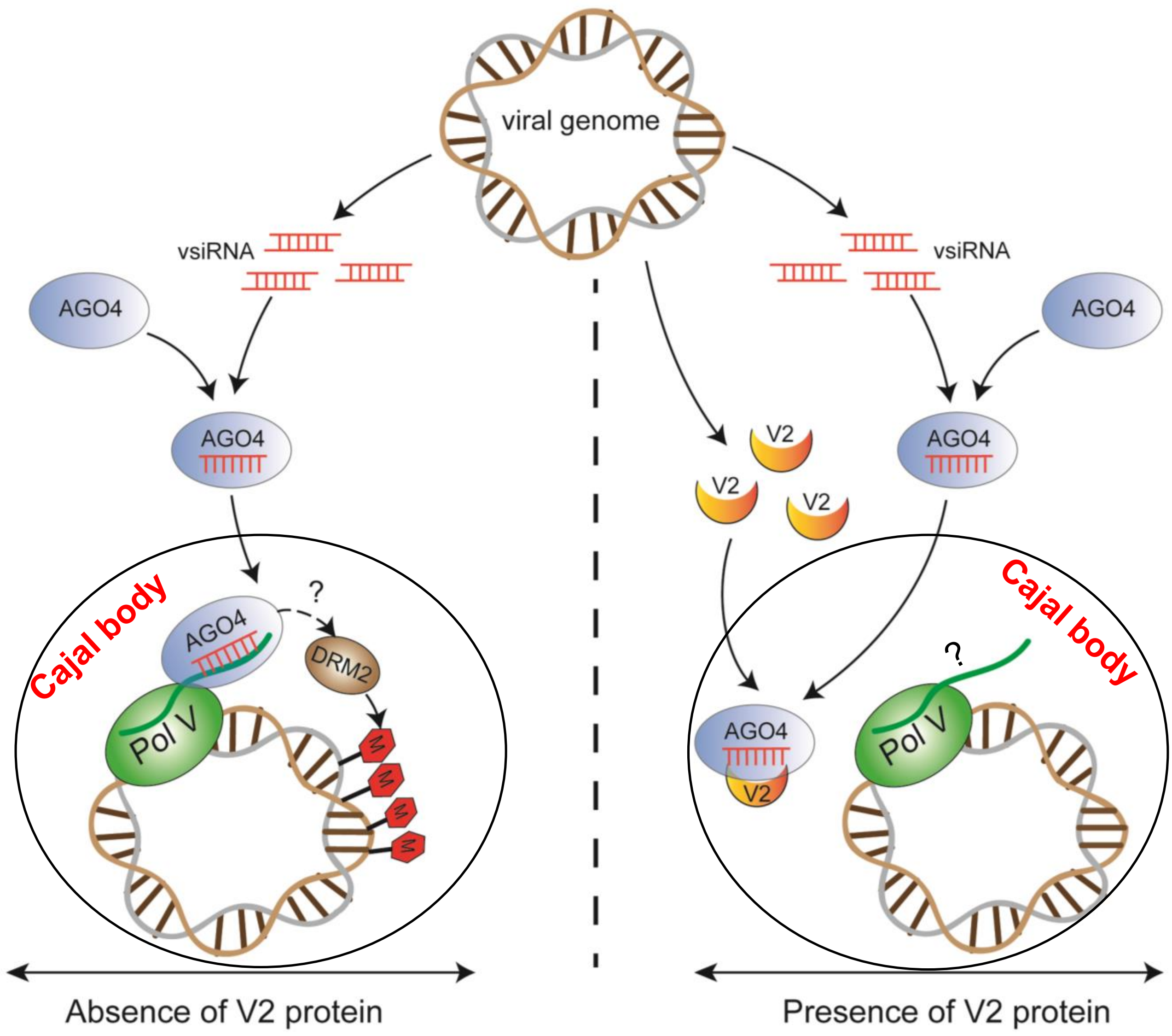


**Figure S8. Model for the V2-mediated inhibition of the AGO4-dependent methylation of the viral DNA in the Cajal body.**

During the viral infection, the ssDNA TYLCV genome forms dsDNA replicative intermediates, which could be targeted by the host AGO4-dependent RNA-directed DNA methylation (RdDM) pathway in the Cajal body as an antiviral defence mechanism. Viral small interfering RNA (vsiRNA) are generated and loaded into AGO4. In the absence of the virus-encoded V2 protein, the AGO4-vsiRNA complex could be effectively guided towards the viral genome by complementary base pairing to the scaffold RNA and association with Pol V, and recruit the methyl transferase DRM2 to catalyze methylation of the viral genome. When V2 is present, however, V2 interacts with AGO4 and interferes with the binding of this protein to the viral DNA, enabling viral evasion from the AGO4-dependent DNA methylation.
